# An unexpected bifurcation in the Pointed transcriptional effector network contributes specificity and robustness to retinal cell fate acquisition

**DOI:** 10.1101/2020.08.03.235184

**Authors:** Chudong Wu, Jean-François Boisclair Lachance, Michael Z Ludwig, Ilaria Rebay

## Abstract

Spatiotemporally specific and robust cell fate transitions are fundamental to the development of appropriately patterned tissues. In the *Drosophila* retina, receptor tyrosine kinase / mitogen activated protein kinase (MAPK) signaling acts through the transcriptional effector Pointed (Pnt) to direct two distinct rounds of photoreceptor specification. A relay mechanism between two Pnt isoforms, a MAPK responsive form PntP2 and a constitutively active form PntP1, initiates and sustains the transcriptional response. Here, we report an unexpected bifurcation in the Pnt effector network. We show that PntP2 works redundantly with a closely related but previously uncharacterized isoform, PntP3, to activate *pntP1* during specification of first round photoreceptors. Intrinsic activity differences between PntP2 and PntP3, combined with positive and negative transcriptional auto- and cross-regulation, buffer first-round fates against conditions of low signaling. In contrast, in a mechanism that may be adaptive to the stronger signaling environment used to specify second round fates, PntP2 uniquely activates *pntP1*. We propose that differences in expression patterns, transcriptional activities and regulatory interactions between Pnt isoforms together facilitate context-appropriate cell fate specification in different signaling environments.

## Introduction

During development, cells integrate external signals and internal information to coordinate the transition from a multipotent to a differentiated state. A relatively small number of transcription factors acting downstream of an even smaller handful of signal transduction pathways coordinate the gene expression changes that drive cell fate acquisition (Flores et al., 2000; Halfon et al., 2000; Voas and Rebay, 2004). Two conditions must be fulfilled to effect these transitions: specificity, whereby cells adopt the correct fate in a precise spatiotemporal manner (Cagan, 2009; Guillemot, 2007; Wolpert, 1969); and robustness, whereby cells reliably execute the appropriate program despite genetic and nongenetic variations (Félix and Barkoulas, 2012; Liu et al., 2019). Despite recent progress, the regulatory strategies used by transcription factors to elicit specific and reproducible developmental transitions remain poorly understood.

Photoreceptor specification during Drosophila retinal development offers an ideal model to study the regulatory mechanisms that confer specificity and robustness to cell fate acquisition. Each of the ~750 ommatidia that comprise the compound eye contains a core cluster of eight photoreceptors, R1-R8. These neurons are specified in a stereotyped spatiotemporal sequence that is initiated repeatedly as the morphogenetic furrow (MF) travels anteriorly across the epithelial field (Wolff and Ready, 1991). Photoreceptor specification occurs in two rounds that are separated temporally by a single synchronized cell division known as the second mitotic wave (SMW) (Ready and Hanson, 1976; Tomlinson and Ready, 1987). During the first round, R8, the founder cell of each ommatidium, emerges from the MF’s wake, followed by the R2/R5 and R3/R4 pairs. During the second round, specification of photoreceptors R1/R6 and finally R7 completes the cluster. Recruitment of non-neuronal support cells to the ommatidia follows immediately, starting with the four lens-secreting cone cells.

Specification of all photoreceptors except R8 requires inductive signaling by the receptor tyrosine kinase (RTK) / Ras / mitogen-activated protein kinases (MAPK) pathway via the transcriptional effector Pointed (Pnt), the Drosophila homologue of the mammalian ETS family activators ETS1 and ETS2 (Freeman, 1996; Scholz et al., 1993). Multipotent retinal progenitors must therefore translate this common signal into specific photoreceptor fates. Numerous studies have emphasized the importance of combinatorial regulation at target gene enhancers as a strategy to integrate generic inputs from RTK/Pnt with specific inputs from regionally expressed transcription factors and other signaling pathway effectors. For example, RTK/Pnt, the Spalt transcription factors and Notch signaling collectively specify R4 fates in the first round (Domingos et al., 2004; Weber et al., 2008) whereas in the second round, RTK/Pnt and Notch signaling integrate with a different transcription factor, Lozenge, to regulate *prospero* transcription and R7 fates (Hayashi et al., 2008; Xu et al., 2000).

Increasing the complexity of these combinatorial codes, RTK signaling inputs are not identical during the two rounds of specification. Fate specification in the first round relies exclusively on signaling initiated by the epidermal growth factor receptor (EGFR); however specification of R1, R6 and R7 second round fates involves a second RTK, Sevenless (Sev) in addition to EGFR (Basler and Hafen, 1988; Reinke and Zipursky, 1988). Although only R7 fates are lost in a *sev* mutant, the R1 and R6 precursors express Sev, physically contact the Boss ligand-expressing R8 cell, and so are likely to have active Sev signaling (Tomlinson et al., 2011). Because EGFR and Sev use the same Ras/MAPK/Pnt signaling cascade, they are considered biochemically interchangeable; indeed, increasing EGFR signaling activity is sufficient to compensate for loss of Sev signaling in R7 fate specification (Freeman, 1996). Given the additional source of MAPK activity conferred from Sev, it has been proposed that cells specified the second round experience stronger MAPK activation than those the first round (Freeman, 1996; Tomlinson et al., 2019). How the Pnt response is tailored to these two different signaling environments to initiate distinct cell fates has not been explored.

Two Pnt isoforms, PntP1 and PntP2, were identified when the gene was first cloned and have been the focus of subsequent study. Use of distinct transcription start sites together with unique 5’ but common 3’ exons produces two proteins with distinct N-terminal transactivating domains, a common C-terminal ETS DNA binding domain (Fig. 1A, B) and unique, non-overlapping expression in some contexts and co-expression in others, including the developing retina (Brunner et al., 1994a; Gabay et al., 1996; Klambt, 1993; O’Neill et al., 1994; Scholz et al., 1993). A key functional distinction between PntP1 and PntP2 is their differential regulation by RTK/MAPK signaling (Fig. 1B, C). Whereas MAPK activation promotes the transcription of *pntP1* (Shwartz et al., 2013) to produce the constitutively active PntP1 transcription factor, MAPK regulation of PntP2 occurs post-translationally by direct phosphorylation of a site within its unique N-terminal half (Brunner et al., 1994a; O’Neill et al., 1994). Thus unphosphorylated PntP2 has limited transcriptional activity and requires phosphorylation by MAPK for full activity.

**Fig. 1.**
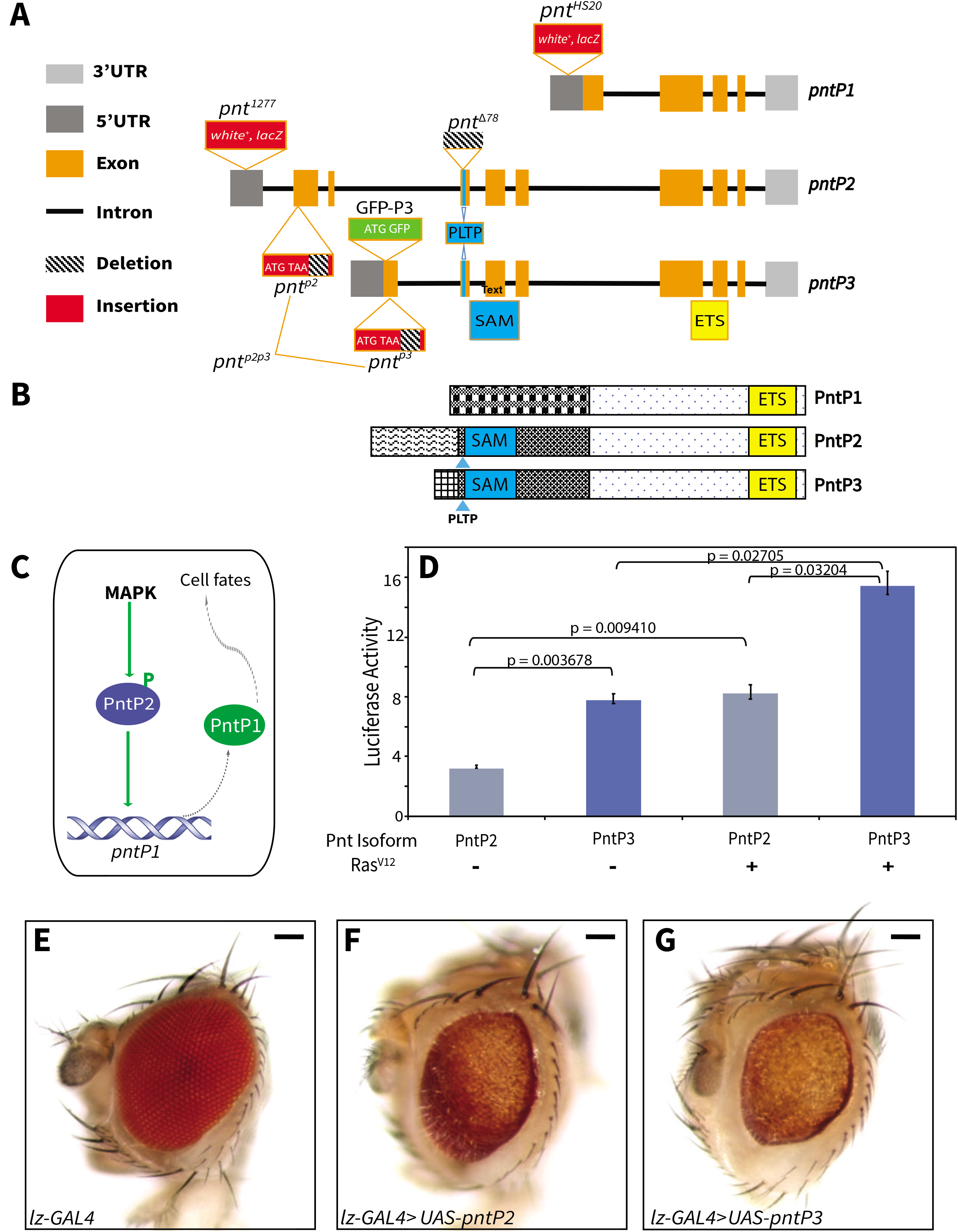
PntP3 is a stronger MAPK-responsive transcriptional activator than PntP2. **(A)** A schematic, not drawn to scale, of the ~55kb *pnt* locus, showing the 5’ and 3’ UTRs, exons and introns of *pntP1, pntP2* and *pntP3*. All three isoforms splice into common 3’ exons encoding the ETS DNA binding domain (yellow box). *pntP2* and *pntP3* also share three internal exons encoding the SAM and PLTP MAPK phosphorylation site (blue boxes). Unique N-terminal exons encode isoform-specific sequences. Approximate insertion sites of key P-element-derived alleles are shown: the *white+, lacZ* enhancer trap insertions *pnt^1277^* and *pnt^HS20^* respectively report *pntP2* and *pntP1* expression (Scholz et al., 1993; Shwartz et al., 2013); the excision allele *pnt^Δ78^* disrupts the SAM-encoding exon common to *pntP2* and *pntP3* (O’Neill et al., 1994). Green boxes labeled ATG-GFP signify genomic BAC transgenes in which PntP3 was N-terminally GFP tagged. Red boxes labeled ATG-TAA represent the CRISPR-generated null alleles of *pnt^p2^* and *pnt^p3^* in which stop codons were introduced immediately after the ATG; both alleles also carry exonic deletions (see Methods); *pnt^p2p3^* carries identical stop codon insertions and deletions. **(B)** A schematic of PntP1, PntP2 and PntP3 highlighting their distinct N-termini and common C-termini. PntP2 and PntP3 differ only in the short stretch of sequence immediately N-terminal to the MAPK site and SAM (141aa for PntP2 and 59aa for PntP3). **(C)** A schematic summarizing the sequential activation model: MAPK signaling activates PntP2 by direct phosphorylation, phosphorylated PntP2 activates *pntP1* transcription and PntP1 protein drives cell fate specification (Shwartz et al., 2013). **(D)** PntP3 has stronger activity but similar MAPK responsiveness relative to PntP2 in transcription assays using a reporter with 6 tandem high-affinity ETS sites (O’Neill et al., 1994), show. For each sample, activity was normalized to reporter alone control (see Methods). Error bars are S.D. of three independent experiments. P-values were calculated using two tailed pairwise Student T-tests between the samples indicated. **(E-G)** *lz-GAL4-driven* overexpression of *UAS-pntP2* (**F)** and *UAS-pntP3* **(G)** disrupts external eye morphology, pigmentation and size relative to driver alone control **(E).** Scale bar: 50 μm.

A previous sequential activation model posited that transient RTK/MAPK signaling activates PntP2, which in turn activates *pntP1* transcription, and that PntP1 then provides a stable, signaling-independent, transcriptional input to the combinatorial codes that initiate the specification of R1-R7 photoreceptor fates (Fig. 1C; Shwartz et al. 2013). However the expression patterns of PntP1 and PntP2 suggest further complexity, with high *pntP1* and low *pntP2* in the region of R2-R5 specification and then high *pntP2* and low *pntP1* in more posterior regions where R1,R6, R7 and the cone fates are recruited (Shwartz et al., 2013). The regional complementarity in isoform-specific expression levels motivated us to consider the possibility that the Pnt response might be different during the two rounds of photoreceptor specification.

In this study, we characterize a novel Pnt isoform, PntP3, and explore its contributions to the Pnt regulatory network. We demonstrate that like PntP2, PntP3 functions as a MAPK-responsive transcriptional activator, but with intrinsically higher activity. Using Crispr/Cas9 engineered isoform-specific mutations and recombineered BAC transgenes with fluorescent protein tags, we show that PntP3 and PntP2 are coexpressed and work redundantly to specify photoreceptors R2-R5 whereas PntP2 is uniquely expressed and required for R1, R6 and R7 fates. Mechanistically, we uncover distinct auto- and cross-regulatory transcriptional interactions between *pnt* isoforms during the two rounds of photoreceptor specification. Extension of the analysis to the developing wing emphasizes the potential for both synergistic and unique contributions of PntP2 and PntP3 during patterning. We propose that a combination of functional redundancy, synergistic activity and transcriptional regulatory interactions between Pnt isoforms contributes both specificity and robustness to cell fate transitions in Drosophila.

## Materials and Methods

### Drosophila strains

From the Bloomington *Drosophila* Stock Center: *Lz-Gal4, pnt^Δ88^, pnt^1277^, FRT82b, egfr^co^*. Additional strains: *ro-GAL4* (Mavromatakis and Tomlinson, 2013), *Sev-yan^ACT^* (Rebay et al., 2000), *rl^S135^* (Karim et al., 1996), *ey-FLP; act-Gal4, UAS-GFP/CyO; FRT82b, tub-GAL80/TM6B* (a gift of Wei Du, University of Chicago, IL, USA), *UAS-flag-pntP2, UAS-flag-pntP3*, GFP-PntP3, *pnt^p2^, pnt^p3^, pnt^p2p3^* (this work). For further details of *Drosophila* strains and genetics, see the supplementary Materials and Methods.

### Transcription assays

2.25 × 10^6^ *Drosophila* S2 cells were transfected with 100 ng of 6X-ETS luciferase reporter construct, 100 ng of PntP2/pMTHA or 100 ng of PntP3/pMTHA, 20 ng of actin >Renilla luciferase, and if applicable, 5 ng of Ras^V12^/pMT. For further details of transcription assay experiments, see the supplementary Materials and Methods.

### Immunohistochemistry and microscopy

Eye-antennal and wing imaginal discs were fixed in 4% paraformaldehyde (PFA) for 10 min. For adult retina staining, decapitated adult heads were fixed in 4% PFA for 20 min, and then the dissected retinas were post-fixed in 4% PFA for 10 minutes. Primary and secondary antibodies were incubated overnight at 4°C. For antibody details, see supplementary Materials and Methods. Imaging was performed with a Zeiss LSM 880 confocal microscope, using 0.8 to 1.0 μm steps and projecting maximally through the desired tissue unless otherwise noted. To image adult eyes and wings, decapitated heads and dissected wings were imaged with a Canon EOS Rebel camera fitted to a Leica stereo microscope. Individual slices were merged using iSolution-Lite software (IMT-Digital).

### Quantitative reverse transcription PCR (RT-qPCR)

Total RNA was extracted from 60 pairs of late 3^rd^ instar eye-antennal or wing discs for cDNA synthesis. qPCR was performed using the iTaq Universal SYBR Green Supermix (Bio-Rad Laboratories) on a 7300 Real-time PCR Machine (Applied Biosystems). Subsequent disassociation analysis was performed with 7300 system software to confirm the sequence specificity of the reaction. For further details of RT-qPCR and primers used, see the supplementary Materials and Methods.

### CRISPR/Cas9-mediated generation of *pnt* mutants

Genomic DNA containing the *pntP2* or *pntP3* specific exon regions was amplified by PCR. After insertion of stop codons and deletion of coding sequence, the fragments were inserted into pHD-Scarless (generated by O’Connor-Giles laboratory, *Drosophila* Genomics Resource Center, 1364; Gratz et al., 2013). The resulting templates were confirmed by sequencing. Guide RNAs were subcloned into the pU6-Bbs1 chiRNA plasmid (Addgene, 45946; Gratz et al., 2013). Each template (300 ng/μL) and the two guide RNAs (75 ng/μL), were injected into a GFP/ RFP-negative *vasa-Cas9* strain (a gift from Rick Fehon). To generate *pnt^p2p3^*, the template and guide RNAs used to generate *pnt^p2^* were injected with *nanos-Cas9* plasmid (a gift from Rick Fehon, (Ren et al., 2013)) into the *pnt^p3^* strain. G_0_ adults were crossed individually to *w^1118^*, and transformants were identified by 3X-Pax-RFP expression in the eyes of the F1 progeny. The 3X-Pax-RFP piggyBac cassette was excised, and RFP-negative progeny were crossed to *TM6B* to establish stocks. The alleles were confirmed by restriction digest and sequencing. For further details and sequences of the CRISPR templates and guide RNAs, see the supplementary Materials and Methods.

### Fluorescence In situ hybridization (FISH)

DNA probes conjugated with NHS ester-ATT0 633 fluorophore were used to target specific exons of *pnt* isoforms. Dissected eye discs from white pre-pupae were fixed in 1% PFA, dehydrated with methanol, and incubated with probes at 62°C. Imaging was performed with a Zeiss LSM 880 confocal microscope, using 0.8 μm steps and projecting maximally through the tissue. Maximum projected images were analyzed with Fiji and Microsoft Excel. In all pair-wise comparison of wild type vs. *pnt* mutant, discs from the two strains were dissected, processed and imaged in parallel. For details of probes used, FISH protocol and image analysis, see the supplementary Materials and Methods.

## Results

### PntP3 is a MAPK-responsive transcriptional activator whose expression partially overlaps that of PntP2

Although the field has focused on the two isoforms identified when the *pnt* gene was first cloned (Brunner et al., 1994; Klambt, 1993; O’Neill et al., 1994; Scholz et al., 1993; Shwartz et al., 2013), the BDGP cDNA sequencing project together with subsequent high-throughput mRNA sequencing has revealed additional transcripts (Fig. S1A; Celniker et al., 2009; Leader et al., 2018; Rubin et al., 2000). First, there is a second *pntP1* isoform (identified as *pnt-E* in G-Browse) identical to “classic” *pntP1* (*pnt-C*) except for a longer 3’UTR and an encoded product with an extra two amino acids owing to the use of an alternate splice donor site at the 3’ end of the first coding exon. Second, there is a transcript identified as *pnt-D* in G-Browse that is closely related to but distinct from *pntP2* (*pnt-B*); we refer to this novel isoform as *pntP3*.

*pntP3* is distinguished from *pntP2* by its unique transcription start site, 5’UTR and N-terminal coding exons, but then like *pntP2*, it splices into the exons encoding the sterile alpha motif (SAM) protein-protein binding domain and adjacent MAPK consensus site, and the ETS domain (Fig. 1A, B). To our knowledge there have not been any explicit studies of *pntP3*. However, just like *pntP1* and *pntP2, pntP3* was detected by RNA-Seq profiling in most developmental stages from embryo to adulthood (FlyBase, Leader et al., 2018), suggesting it might contribute to the transcriptional response downstream of receptor tyrosine kinase (RTK) signaling. Further, both PntP2 and PntP3 are conserved across *Drosophila* species from *D. melanogaster* to *D. virilis* (Fig. S1B). The conservation of the two isoforms across millions of years suggests strong evolutionary pressure for keeping both PntP2 and PntP3, implying essential functions.

Given the protein-level similarity, we began by asking whether PntP3, like PntP2, functions as a transcriptional activator positively regulated by MAPK phosphorylation. In transcriptional reporter assays in transiently transfected S2 cells, PntP3 was about two-fold more active than PntP2 in the absence of MAPK stimulation; MAPK stimulation induced a further ~two-fold activity increase for both isoforms (Fig. 1D). When overexpressed in the developing eye, PntP3 also showed greater activity than PntP2, producing stronger disruptions of adult eye morphology with all Gal4 drivers tested (Fig. 1E-G, Fig. S2 and not shown). These phenotypes were associated with ectopic induction of photoreceptor fates in 3^rd^ instar discs, consistent with previous studies of Pointed overexpression (Shwartz et al., 2013). As predicted by their relative activities in S2 cells, ectopic expression of neuronal markers was more striking with *pntP3* overexpression than with *pntP2*, (Fig. S2A-F), The expression level and subcellular localization to the nucleus were indistinguishable between the two isoforms (Fig. S2 G-I), indicating differential transcriptional activity most likely underlies the phenotypic differences. Together these results suggest that PntP2 and PntP3 have similar biochemical function but that the distinct N-terminal sequence of PntP3 confers higher intrinsic transactivation potential.

In addition to the activity differences associated with the unique N-terminal sequences of PntP2 versus PntP3, the use of separate 5’ regulatory regions suggested that expression pattern differences might also distinguish their developmental roles. To explore this possibility, we compared their endogenous expression in late 3^rd^ instar eye discs, where *pnt* function is essential for photoreceptor specification and has been well studied (Brunner et al., 1994a; Scholz et al., 1993; Yang et al., 2003). We relied on the *pntP2*-specific enhancer trap allele *pnt^1277^* (Scholz et al., 1993; Shwartz et al., 2013) to report PntP2 expression, whereas to visualize PntP3 expression, we inserted a GFP tag in-frame at its very N-terminus in a genomic BAC that contains the entire *pnt* locus (GFP-P3, Fig. 1A) and that we had previously shown to be fully functional (Lachance et al., 2014). The *GFP-PntP3* transgene fully complemented the lethality of *pnt^Δ88^/Df(pnt)* animals, a background null for all three isoforms, producing phenotypically wild type, fertile adults.

Analysis of 3^rd^ instar eye discs dissected from animals carrying both *GFP-PntP3* and the *pntP2*-specific enhancer trap allele revealed both distinct and overlapping patterns of expression (Fig. 2). Lower magnification projections emphasized the complementary aspects of the two patterns, with GFP-PntP3 expression strongest in and immediately posterior to the MF and β-galactosidase (β-gal) reporting strongest *pntP2* expression in the posterior half of the eye field (Fig. 2B). As the stereotyped differentiation sequence of ommatidial assembly means every cell can be unambiguously identified by its position and morphology (Ready et al., 1976; Tomlinson et al., 1987; Wolff et al., 1993; Pelaez et al., 2015), higher magnification views at different optical planes enabled cell type specific comparison of the two patterns (Fig. 2C-F). Coexpression was detected in R2/R5 and R3/R4 photoreceptor pairs, in basal progenitors at the MF and in cone cells (Fig. 2C, D and S3A). Complementary expression was detected posterior to the second mitotic wave (SMW) in photoreceptors R1, R6 and R7 where *pntP2* was high and GFP-PntP3 low, and in apically localized nuclei at the MF, including R8, where GFP-PntP3 was high and *pntP2* low (Fig. 2E, F). The combined differences and similarities in cell type specific expression patterns raised the possibility of both distinct and overlapping functional requirements for PntP2 and PntP3 in the two rounds of photoreceptor specification (Fig 2A).

**Fig. 2.**
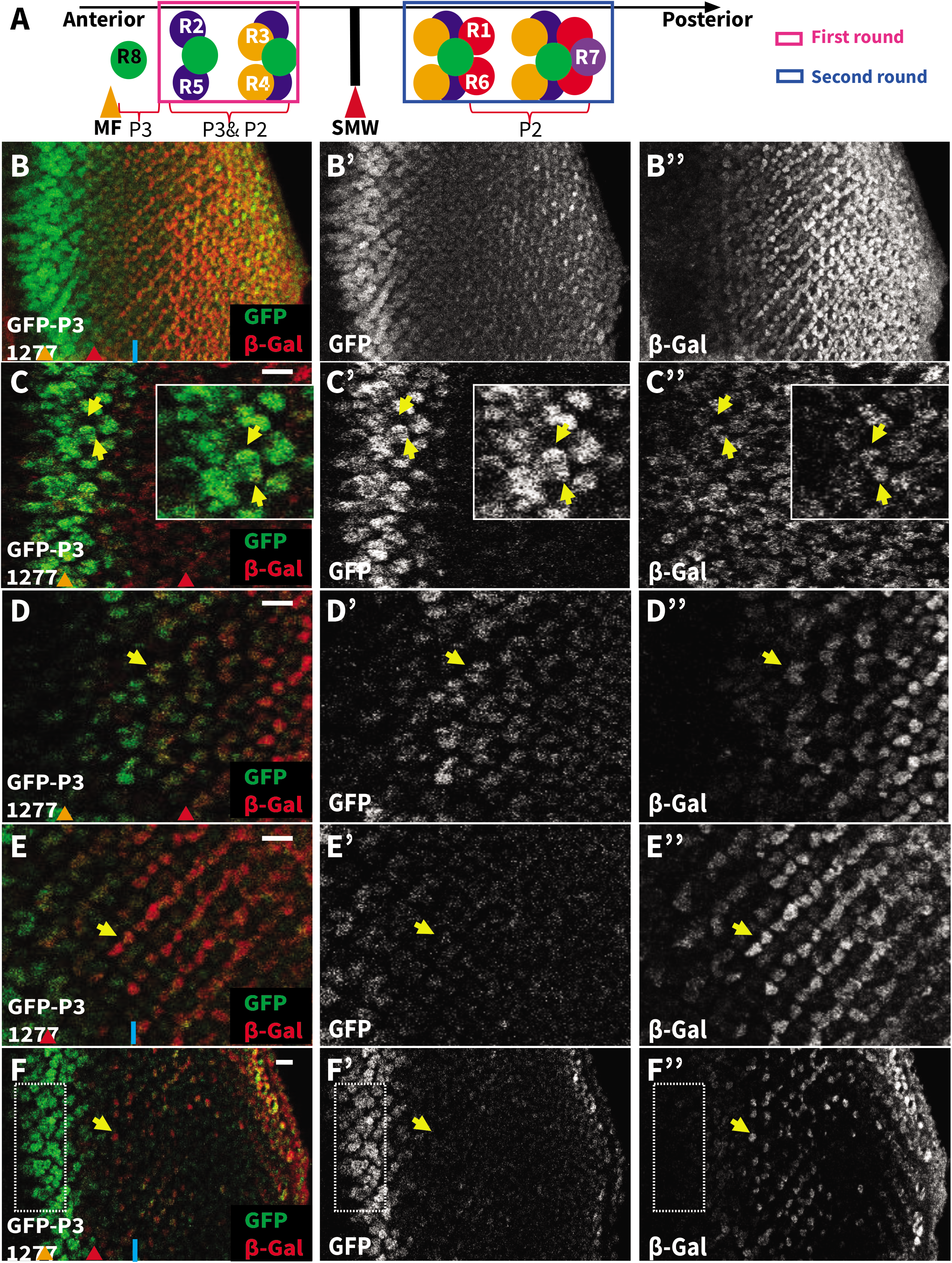
PntP3 and PntP2 show overlapping and complementary expression patterns. **(A)** A schematic summarizing the sequential specification of photoreceptor fates (adapted from (Pelaez et al. 2015)) and the expression patterns of PntP3 and PntP2. R8 cells are specified first near the MF (orange arrowhead) and express PntP3. R2/R5 and R3/R4 pairs are specified next (first round fates) and express both PntP3 and PntP2. After the SMW (red arrowhead), R1/R6 and R7 are specified (second round fates) and express PntP2. **(B-F)** Representative 3rd instar eye imaginal discs, oriented anterior left, comparing the pattern of *pntP2* transcription (red), as reported by *pnt^1277^*, and PntP3 protein (green), as reported by a GFP-PntP3 genomic BAC transgene. MF, orange arrowheads and SMW, red arrowheads. (B) A maximum projection highlights the complementarity in expression, with highest PntP3 in the MF region and highest *pntP2* posterior to the SMW (blue line). (B’, B”) Single channel images show that the onset of *pntP2* transcription occurs anterior to the SMW in cells where GFP-PntP3 is expressed. (C-F) Single optical slices of the disc in (B) at different apical/basal planes. (C, D) Coexpression was detected in R2/R5 pairs (C, yellow arrows, insets show zoomed view) and in R3/R4 pairs (D, yellow arrows). (E-F) *pntP2* but not PntP3 was detected in R1/R6 pairs (E, yellow arrows) and in R7 (F, yellow arrows). PntP3 but not *pntP2* was detected in basal progenitors at the MF (F, boxed region). Scale bar: 10 μm.

### Redundant and non-redundant requirements for PntP2 and PntP3 during two distinct rounds of photoreceptor specification

Prior studies of *pnt* function during retinal development concluded that *pntP1* and *pntP2* are required non-redundantly to specify photoreceptors R1-R7 (O’Neill et al., 1994), providing the foundation for the sequential activation model (Shwartz et al., 2013; and Fig 1C). However the *pntP2* allele used in the studies, *pnt^Δ78^* (O’Neill et al. 1994), was generated by imprecise excision of a P-element inserted into the first SAM-encoding exon, and so also disrupts *pntP3* (Fig. 1A). This means that *pnt^Δ78^* phenotypes, in the eye loss of R1-R7, reflect the combined loss of *pntP2* and *pntP3*.

To reveal the individual requirements for the two isoforms we generated *pnt^p2^* and *pnt^p3^* specific mutants using CRISPR/Cas9 genome editing to insert stop codon(s) right after the start codon of each isoform (Fig. 1A). To confirm the effectiveness of the molecular strategy, we also engineered a *pnt^p2p3^* double mutant allele. As reported for *pnt^Δ78^*(Morimoto et al., 1996; O’Neill et al., 1994), homozygous *pnt^p2p3^* adults were never recovered, indicating that the combined function of the two isoforms is essential for viability. In contrast, homozygous *pnt^p3^* animals were fully viable while homozygous *pnt^p2^* animals occasionally survived. From a cross between balanced heterozygous *pnt^p2^* parents, only 35 homozygous *pnt^p2^* animals were found from a total of 2058 progeny scored at the 3^rd^ larval instar stage; this 1.7% survival rate to 3^rd^ instar is significantly lower than the 33% expected for full viability. The differences in survival of the isoform specific mutants suggested both redundant and non-redundant requirements for PntP2 and PntP3 during development, with PntP2 playing the major role and PntP3 a more auxiliary one.

Focusing on photoreceptor specification, homozygous *pnt^p2p3^* clones were missing all photoreceptors except R8 (Fig. 3A-A’’); this phenotype is consistent with published analysis of *pnt^Δ78^* mutant clones (O’Neill et al., 1994; Shwartz et al., 2013). We reasoned that if the function of both PntP2 and PntP3 is required for photoreceptor specification, then neither single mutant should recapitulate the double mutant phenotype. If so, we predicted the requirement for PntP3 should manifest in the first round fates where it is strongly expressed but not in second round fates where its levels are low (Fig. 2A, B and S3B). Alternatively, if PntP3 does not contribute activity essential to photoreceptor specification, then the *pnt^p2^* and *pnt^p2p3^* mutants should show identical loss of R1-R7 phenotypes.

**Fig. 3.**
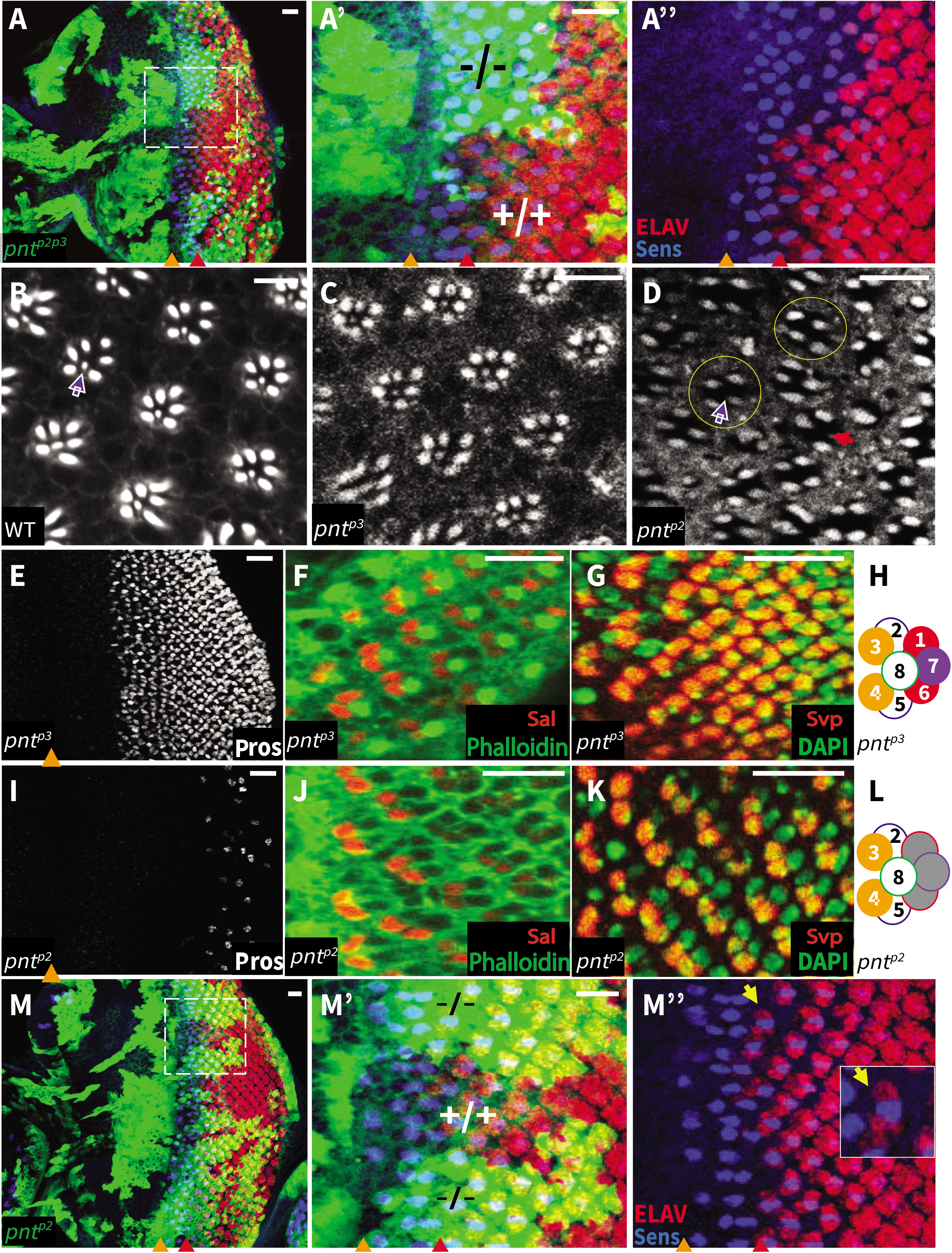
Redundant and unique requirements for PntP2 and PntP3 in photoreceptor specification. **(A)** Representative 3^rd^ instar eye disc of genotype *eyFLP/+; act-Gal4, UAS-GFP; FRT82B, pnt^p2p3^/tub-Gal80, FRT82B*, oriented anterior left, stained with anti Elav (red) to mark all photoreceptors and anti-Sens (blue) to mark R8. Orange arrowheads mark the MF. Homozygous *pnt^p2p3^* mutant clones, positively marked by GFP (green), lack all photoreceptors except R8. A’ and A” show zoomed views of boxed region in A. Scale bar: 10 μm. **(B-D)** Adult retinas stained with phalloidin to mark the actin-rich rhabdomeres. (B, C) Both wild type (WT) and *pnt^p3^* mutants have regularly arrayed rows of ommatidia; white arrow points to the small R7 rhabdomere at the center of the outer trapezoid formed by the larger R1-R6 rhabdomeres. (D) All ommatidia of homozygous *pnt^p2^* mutants lack the R7 rhabdomere, most also lack two outer rhabdomeres (yellow circles), and some show even greater loss (red arrow). Scale bars: 5 μm. **(E-L)** Representative 3rd instar eye imaginal discs, oriented anterior left, with MF marked by orange arrowhead. R7 and cone cells are marked by Pros (white), R3/R4 pairs are marked by Sal (red) or Svp (red), and R1/R6 pairs are marked by Svp (red). (E-G) *pnt^p3^* mutants appear wild type. (I-K) *pnt^p2^* mutants lack R7 and most cone cells (I), have normal R3/R4 specification (J, K), and lack R1/R6 (K). (H, L) Schematic summaries of photoreceptor specification patterns in *pnt^p3^* and *pnt^p2^* mutants. Scale bars: 10 μm. **(M)** Representative 3^rd^ instar eye disc of genotype *eyFLP/+; act-Gal4, UAS-GFP; FRT82B, pnt^p2^/tub-Gal80, FRT82B*, oriented anterior left, stained with anti Elav (red) and anti-Sens (blue). MF and SMW are noted with orange and red arrowheads, respectively. Homozygous *pnt^p2^* clones, positively marked by GFP (green), show normal R8/R2/R5 specification. K’ and K” show zoomed views of boxed region in K; yellow arrow points to a newly specified R2/R5 pair. Scale bar: 10 μm.

To distinguish between these two possibilities, we first assessed photoreceptor loss in adult eyes, using F-actin to highlight the number and spatial arrangement of the rhabdomeres. In a wild type ommatidium, the larger rhabdomeres of R1-R6 are arrayed in a trapezoidal-shaped ring with the smaller rhabdomeres of R7 in its center (Fig. 3B, arrow points to R7). Whereas ommatidia of homozygous *pnt^p3^* adults had the full complement of photoreceptors (Fig. 3C), those of the rare homozygous *pnt^p2^* adults were missing photoreceptors (Fig. 3D). Loss of R7 was fully penetrant (100%, n=204, Fig. 3D, white arrow), with most ommatidia missing two additional photoreceptors (73%, n=204, Fig. 3D, yellow circles), and some showing even greater loss (Fig. 3D, red arrow). The more modest photoreceptor loss seen in *pnt^p2^* single mutants (Fig. 3D) relative to *pnt^p2p3^* double mutants (Fig. 3A) indicates a functional requirement for PntP3.

As an independent test of this conclusion, we crossed our new *pnt* alleles to flies carrying a *Sev-Yan^ACT^* transgene, a genetic background in which constitutive activity of the RTK antagonist Yan blocks specification of the photoreceptor fates in which it is expressed (Rebay et al., 1995). This background had been shown previously to be sensitive to *pnt* dose such that *pnt* heterozygosity dominantly enhanced the *Sev-Yan^ACT^* photoreceptor loss phenotype (Rebay et al., 2000). Removal of one copy of either *pnt^p2^* or *pnt^p3^* dominantly enhanced the *Sev-Yan^ACT^* rough eye phenotype and loss of photoreceptors, while loss of either both copies of *pnt^p3^* or one copy each of *pnt^p2^* and *pnt^p3^* enhanced even further (Fig. S4A-G). Quantification of photoreceptor numbers showed that homozygous loss of *pnt^p3^* and heterozygous loss of *pnt^p2p3^* were phenotypically equivalent with respect to *Sev-Yan^ACT^* enhancement (Fig. S4G). Thus, PntP3 contributes to photoreceptor specification, with redundancy between PntP2 and PntP3 buffering completely against PntP3 loss and partially against PntP2 loss.

In contrast, and consistent with expression pattern-based predictions, loss of *pnt^p2^* resulted in loss of cell fates recruited during the second round of specification. First, only a few Pros positive cells remained in the posterior of discs from homozygous *pnt^p2^* animals; a similar posterior scattering of Cut-positive cells suggested a complete failure to specify R7 photoreceptors and most cone cells (Fig. 3I and Fig. S4H, I). Second, normal expression of Sal and reduction of Svp expression to only two, rather than four cells per ommatidia, indicated correct specification of photoreceptors R3/R4 and a failure to specify R1/R6 (Fig. 3J-L). Third, and confirming no other consistent photoreceptor specification defects, examination of Sens and Elav patterns in *pnt^p2^* mosaic discs indicated normal recruitment of photoreceptors R8/R2/R5 (Fig. 3M, L). Thus the complete loss of R1-R7 fates that occurs in *pnt^p2p3^* double mutant ommatidia (Fig. 3A) reflects the combined loss of redundant inputs to R2-R5 first found fates plus the PntP2-specific input to R1, R6 and R7 second round fates.

### Synergistic and unique functions of Pnt isoforms during wing patterning

PntP2 has been implicated in EGFR-mediated regulation of wing disc patterning (Paul et al., 2013). EGFR signaling initially specifies the dorsal compartment and then as the disc grows, continued strong signaling specifies the notum, restricting development of the wing proper to the rest of the epithelium (Campbell, 2000; Pallavi, 2003; Paul et al., 2013; Zecca and Struhl, 2002). Reflecting these roles, partially duplicated wing pouches were reported in hypomorphic *pntP2*-specific allelic combinations (Scholz et al., 1993). As predicted by this prior work, homozygous *pnt^p2^* null mutant adult escapers had duplicated and malformed wings (Fig. 4B, green line marks the wing tissue); examination of 3^rd^ instar wing discs revealed duplication of the wing pouch and reduction of notum tissue (Fig. 4A, C, yellow arrow). In contrast, *pnt^p3^* null mutant wings and wing discs were normally patterned (Fig. 4D and not shown).

**Fig. 4.**
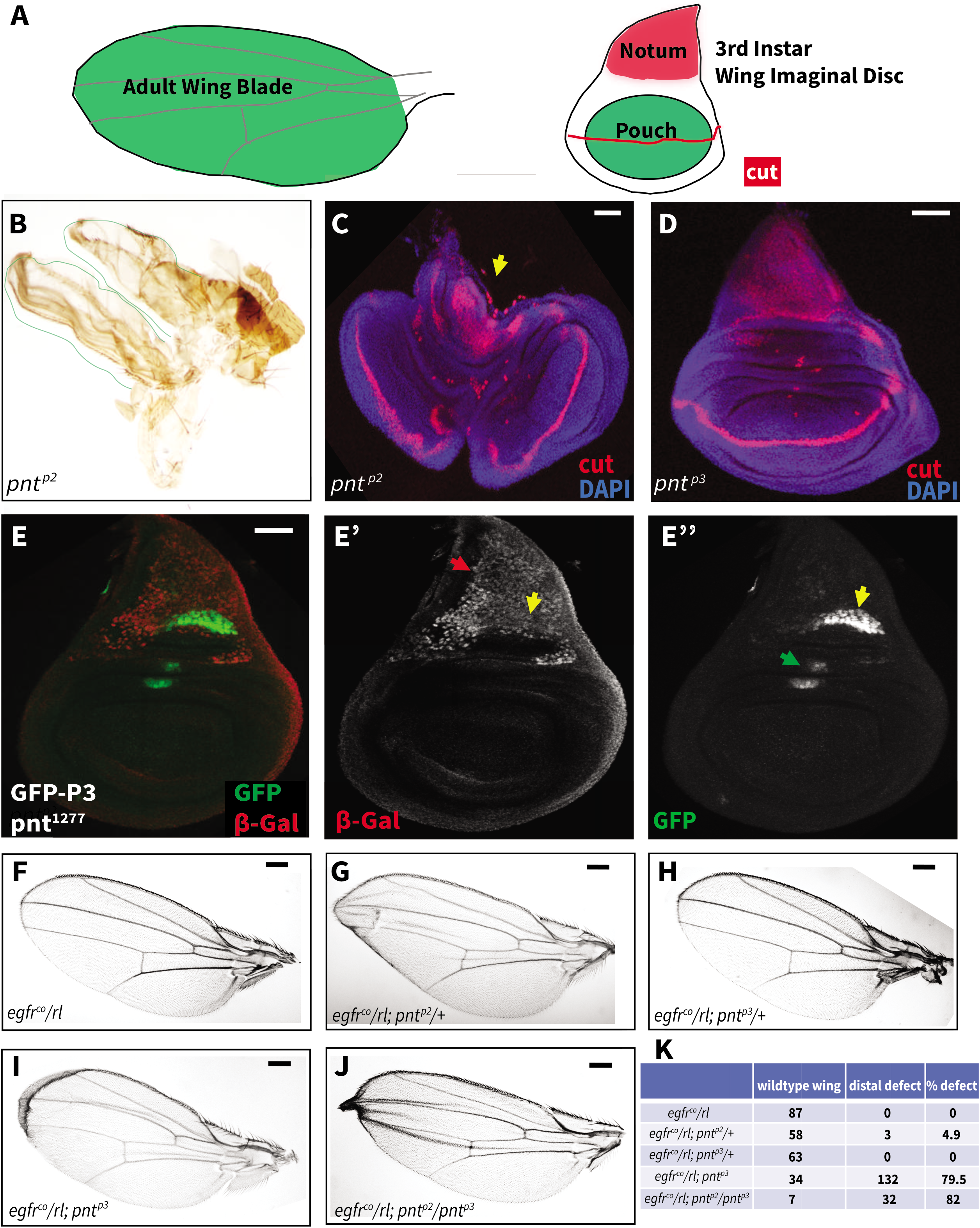
The unique and synergistic functions of PntP2 and PntP3 during wing patterning. Adult wings are oriented anterior up and distal left. Maximum projections of 3^rd^ instar wing imaginal discs are oriented dorsal up. **(A)** A schematic depicting the adult wing blade that develops from the pouch region (green) of the wing imaginal disc. Cut expression (red) marks the prospective wing margin. **(B-C)** Loss of *pnt^p2^* results in duplication of wing material. (B) A duplicated and malformed wing dissected from an adult (B) or 3^rd^ instar (C) *pnt^p2^* escaper. Cut expression (red) highlights the wing pouch duplication and reduced notum (yellow arrow) of *pnt^p2^* discs (C) and the wild type patterning of *pnt^p3^* discs (D). DAPI (blue) marks all nuclei. Scale bar: 50 μm. **(E)**β-gal expression from the *pnt^1277^* (red) is seen throughout the notum (red arrow) and overlaps GFP-PntP3 (green) in the posterior compartment (yellow arrow). PntP3 is also expressed in the dorsal hinge region (green arrow). No expression of either isoform was detected in the pouch. Scale bar: 50 μm. **(F-K)** Representative adult wings showing the effects of reduced *pnt^p2^* and *pnt^p3^* dose in a sensitized *egfr^co^/rl* background. (F) *egfr^co^/rl* trans-heterozygotes appeared wild type. (G) loss of one copy of *pnt^p2^* produced occasional distal margin defects. (H) loss of one copy of *pnt^p3^* did not produce patterning defects. (I) loss of both copies of *pnt^p3^* resulted in distal margin defects in ~80% of wings. (J) Simultaneous reduction in dose of both *pnt^p2^* and *pnt^p3^* produced a synergistic increase in wing margin defects. (K) Quantification scoring wings as either wild type or having distal margin defects for each genotype in (F-J). Scale bar: 0.1 mm.

To explore whether redundancy with PntP2 might mask the role of PntP3 in the wing, we first compared their expression patterns in the 3^rd^ instar wing disc. As in the eye, both overlapping and non-overlapping expression domains were detected (Fig. 4E). Thus both isoforms were expressed in a cluster of cells in the posterior compartment of the notum (Fig. 4E’, E”, yellow arrow), cells elsewhere in the notum expressed primarily *pntP2* (Fig. 4E’, red arrow), cell clusters in the dorsal hinge region expressed primarily GFP-PntP3 (Fig. 4E”, green arrow), and neither showed strong expression in the wing pouch.

Because the regional specificity of the expression domains made deducing PntP3 function by comparing the phenotypes of *pnt^p2^* versus *pnt^p2p3^* somatic mosaics difficult, we instead assessed the consequences of reducing *pntP3* dose in a sensitized genetic background in which animals were doubly heterozygous for null alleles of *egfr* and *rolled*(*rl*). The wings of the double heterozygotes were wild type, indicating that EGFR signaling remained adequate to support normal development (Fig. 4F). Confirming the background was indeed sensitized, removal of one copy of *pntP2* produced defects in the distal wing margin and creases along the longitudinal axis, albeit at low penetrance (Fig 4G, K). Although heterozygosity for *pntP3* was not sufficient to produce a phenotype in the *egfr/rl* background (Fig. 4H, K), when both copies were removed, 80% of the adults showed distal wing margin defects and longitudinal creases (Fig. 4I, K). Margin defects and creases also occurred in *egfr/rl; pnt^p2^/pnt^p3^* quadruple heterozygotes (Fig. 4J); the 82% penetrance of these phenotypes relative to the low penetrance in the single heterozygotes (Fig 4K) suggested strong synergy between *pntP2* and *pntP3*. Given that neither isoform had detectable expression in the third instar wing pouch, these adult phenotypes must reflect loss of expression at a different stage. Further work will be required to identify when and where PntP2 and PntP3 are coexpressed in the wing pouch in order to understand better these phenotypes.

### PntP2 and PntP3 provide robustness through redundant activation of pntP1 transcription

A central tenet of the current model of *pnt* function during photoreceptor specification is that PntP2 activates *pntP1* transcription (Shwartz et al. 2013; Fig. 1C). Given the partial genetic redundancy between PntP2 and PntP3, we asked whether PntP3 also contributes to this activation. To start, we used reverse transcription quantitative polymerase chain reaction (RT-qPCR) to measure *pntP1* transcript levels in *pnt^p2^* and *pnt^p3^* mutant tissues. In both mutants, modest, but not significant decreases in *pntP1* transcripts were measured in eye discs (p = 0.1; Fig. 5A) and no changes were detected in wing discs (Fig. S5A). This suggests either redundancy between PntP2 and PntP3 with respect to activating *pntP1* transcription, inadequate sensitivity in the RT-qPCR assay, or that PntP2 and PntP3 are not the primary activators of *pntP1*.

**Fig. 5.**
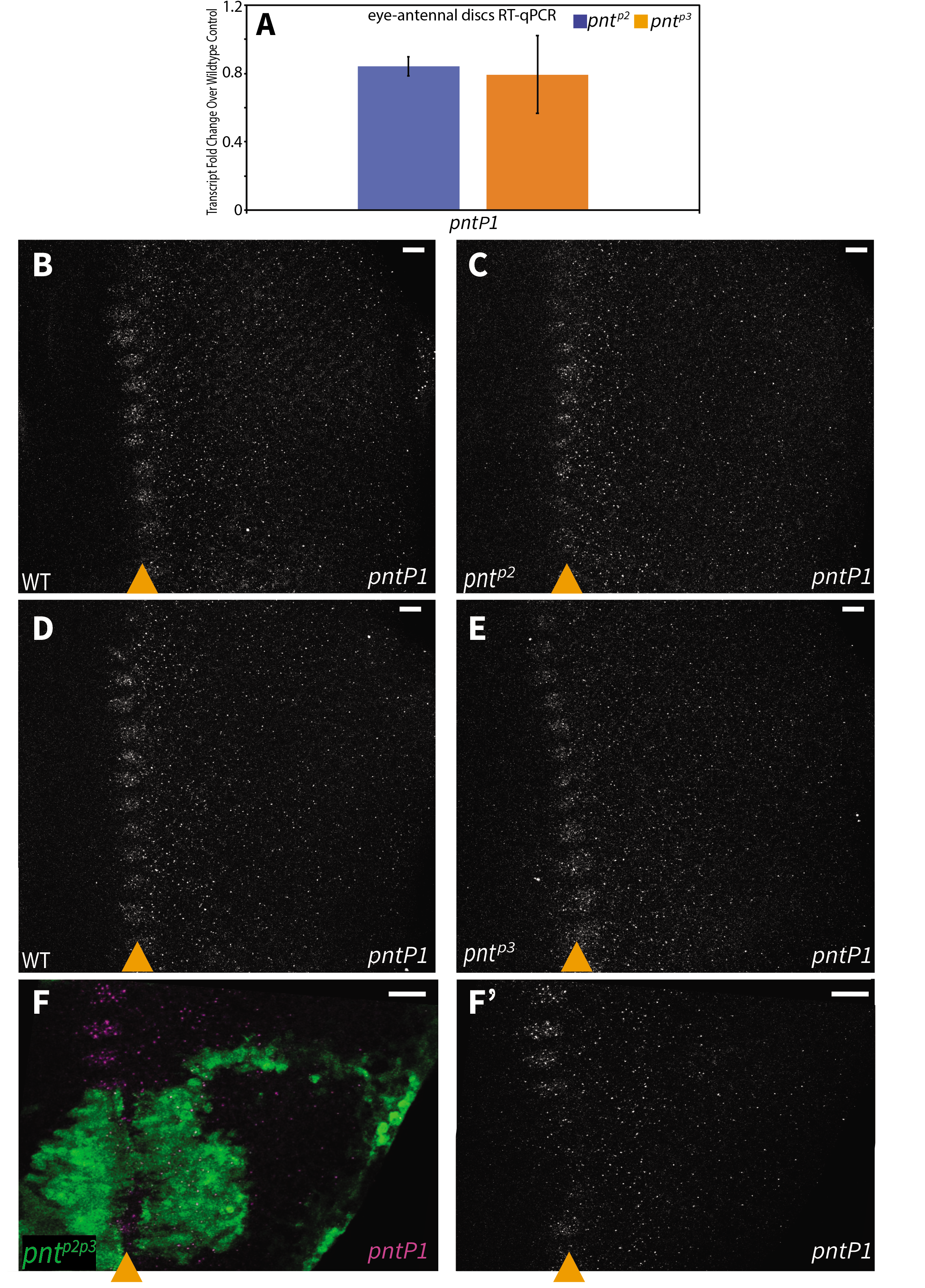
PntP2 and PntP3 redundantly activate *pntP1* transcription. **(A)** RT-qPCR comparison of *pntP1 transcript* levels in wild type versus *pnt^p2^* (blue bars) and *pnt^p3^* (orange bars) 3^rd^ instar eye-antennal imaginal discs. No significant changes were detected. Error bars represent S.D. of three independent experiments. Significance was calculated via pairwise Student T-tests between the mutant sample and the control gene. **(B-F)** *pntP1* FISH in 3^rd^ instar eye imaginal discs, oriented anterior left, with MF marked by orange arrowhead. (B-E) Maximum projections, (F) partial projections of the representative clone regions. Scale bars: 5 μm. **(B, C)** *pntP1* transcripts were detected in a periodic pattern at the MF and then at a low uniform level across the wild type eye field (B). No obvious changes were detected in *pnt^p2^* discs (C). **(D, E)** *pntP1* transcript patterns were comparable between wild type (D) and *pnt^p3^* (E). **(F)** Homozygous *pnt^p2p3^* mutant clones, positively marked with GFP (green), show reduced *pntP1* levels and loss of the periodic pattern at the MF. Consistent results were obtained from analyzing 12 clones from 9 discs across 3 independent experiments.

We were concerned that by grinding up whole tissue we were destroying spatial information and therefore missing locally significant changes in *pntP1* levels. Also, the animals lacking both PntP2 and PntP3 do not survive to 3^rd^ instar, precluding RT-qPCR analysis of the double mutant. Thus we turned to fluorescence in situ hybridization (FISH) to ask whether the two isoforms work redundantly to active *pntP1* expression. To start we examined *pntP1* transcription, using probes that target its isoform-specific exons. The FISH showed an expression pattern consistent with that of the *pntP1* enhancer trap allele (Scholz et al., 1993; Shwartz et al., 2013). Specifically, we detected peak *pntP1* transcription in a periodic pattern at the MF, lower levels of expression extending to the SMW region and then decreased levels in the posterior of the disc (Fig. 5B, D). In pair-wise comparisons of wild type vs. *pnt^p2^* and wild type vs. *pnt^p3^*, no changes in *pntP1* transcription were noted (Fig. 5B-E), consistent with the RTqPCR results. However in *pnt^p2p3^* mutant clones, *pntP1* expression at the MF was strongly reduced (Fig. 5F-F’). We conclude that PntP2 and PntP3 redundantly activate *pntP1*.

### Context specific auto- and cross-regulation of *pntP2* transcription

Having established the functional redundancy of PntP2 and PntP3 with respect to induction of *pntP1*, we next investigated how the system tunes these two parallel inputs to achieve the desired output. In particular we wondered whether cross-regulatory feedback might coordinate and optimize PntP2/PntP3 expression levels, and ultimately their activity. To test this possibility, we used RT-qPCR to measure changes in *pntP2* and *pntP3* transcript levels in eye imaginal discs dissected from *pnt^p2^* and *pnt^p3^* homozygous mutant 3^rd^ instar larvae.

Two findings emerged. Most striking, and unexpectedly, the experiment uncovered negative auto-regulation for both isoforms (Fig. 6A). Thus, *pntP2* transcript levels were significantly increased in *pnt^p2^* mutant tissue (p < 0.01) and *pntP3* transcripts were significantly increased in *pnt^p3^* mutant tissue (p < 0.05). Given the surprising nature of this result, we repeated the experiment using wing imaginal discs, and again found significant increases in transcript levels in the respective mutant (Fig. 6B). This suggests that both isoforms negatively regulate their own transcription, either directly or indirectly.

**Fig. 6.**
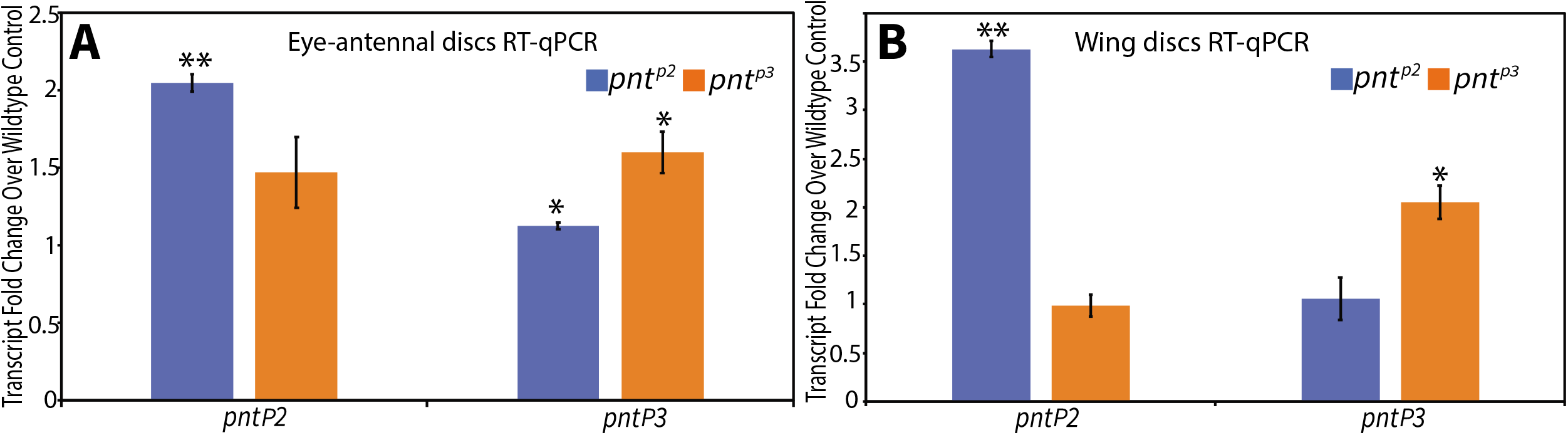
Auto-repression of *pntP2* and *pntP3*. **(A, B)** RT-qPCR comparison of *pntP2* and *pntP3* transcript levels in wild type versus *pnt^p2^* (blue bars) and *pnt^p3^* (orange bars) 3^rd^ instar eye-antennal (A) and wing (B) imaginal discs. Significant increases were detected in both tissues. Error bars represent S.D. of three independent experiments. Significance was calculated via pair-wise Student T-tests between the mutant sample and the control. **, p < 0.01; *, p< 0.05.

Second, evidence of cross-regulation emerged from the eye disc experiments (Fig. 6A), with a significant increase in *pntP3* transcript levels measured in *pnt^p2^* mutant tissue (p < 0.05). A suggestive, but not statistically significant, increase in *pntP2* transcript levels was noted in *pnt^p3^* mutant tissue (p = 0.18), hinting at bidirectional inhibitory cross-regulation. Cross-regulatory interactions were not detected in the wing disc (Fig. 6B).

The coexpression of PntP2 and PntP3 in the anterior half of the disc where first round photoreceptor fates are specified predicted that the regulatory interactions uncovered by RT-qPCR were occurring in this context. We therefore turned to FISH to corroborate the negative auto-regulation and to assess further the possibility of cross-regulatory interactions. We found that *pntP2* transcription initiated at the MF, peaked in the region of the second mitotic wave (SMW), and then continued at a more moderate level across the posterior half of the disc (Fig. 7A, C). This pattern was consistent with that reported by the enhancer trap *pnt^1277^* although the prolonged perdurance of beta-galactosidase likely over-reports *pntP2* levels in the posterior half of the disc (Fig. 2). Unfortunately our FISH protocol was not able to detect *pntP3*, presumably because its specific exon is too short for adequate numbers of probes (see Methods).

**Fig. 7.**
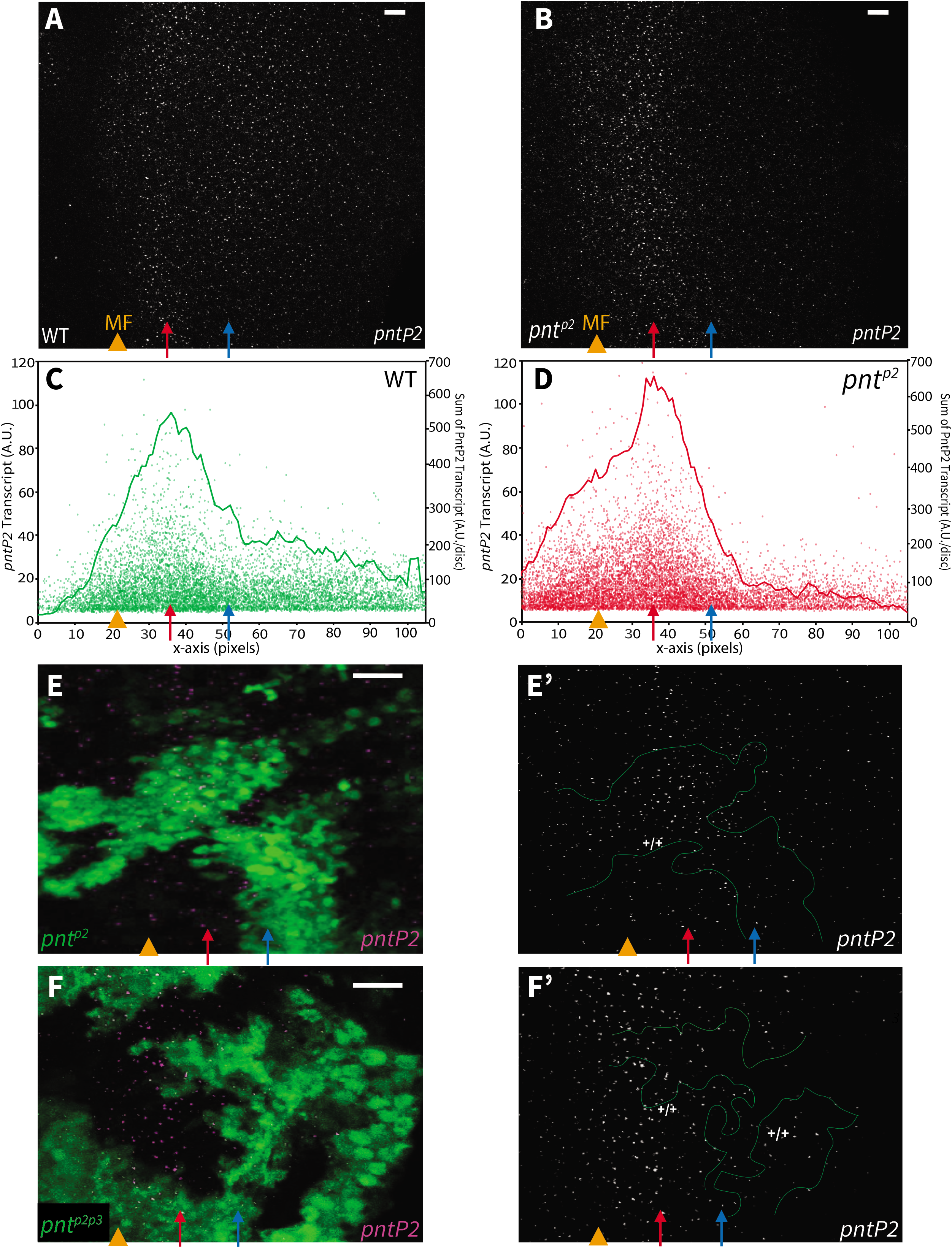
Distinct context-specific interactions regulate *pntP2* transcription across the eye field. **(A-B)** Maximum projection images of *pntP2* FISH in representative wild type and *pnt^p2^* 3^rd^ instar eye imaginal discs, oriented posterior to the right. Orange arrowheads mark the MF, red arrows mark the peak of *pntP2* expression and blue arrows mark the start of lower expression in the posterior half of the disc; the three can be mapped to correspondingly colored marks in Fig. 2A based on the pixel distances. In *pnt^p2^* discs (B) relative to wild type (A), an increased and broader peak of *pntP2* transcripts was detected in and immediately posterior to the MF while a decrease was seen in the posterior half of the disc. Scale bar: 5 μm. **(C-D)** Quantification of *pntP2* FISH in wildtype (C) and *pnt^p2^* mutant (D) from maximum projections of 6 independent discs of each genotype. In wild type *pntP2* levels begin to rise anterior to the MF (orange arrowhead), peak (red arrow) and decrease to a steady state in the posterior half (blue arrow). In *pnt^p2^* discs *pntP2* levels were higher than normal in the anterior half (left of blue arrow) but lower in the posterior (right of blue arrow). Each dot plots the product of the fluorescent intensity and the size of an individual *pntP2* FISH focus, representing the relative amount of *pntP2* transcript (y-axis on the left); one focus was detected per nucleus, with individual foci varying in both size and brightness. The line connects the moving average of the sum of all foci within one-pixel windows along the x-axis (y-axis on the right). Further details in materials and methods. **(E)** Homozygous *pnt^p2^* clones in a 3^rd^ instar eye disc, positively marked with GFP (green). Clone boundary is circled with green line (E’). *pntP2* levels in the mutant clones were increased relative to levels in neighboring wild type tissue in the anterior region (orange arrowhead and red arrow) but appeared decreased in more posterior clones (blue arrow). Examination of 8 clones in 7 discs from 3 independent experiments showed consistent changes. Images are partial projections for representative clone regions, scale bar: 5 μm. **(F)** *pntP2* FISH in homozygous *pnt^p2p3^* clones, positively marked with GFP (green). *pntP2* levels in the mutant clones were indistinguishable from wild type in the anterior regions (orange arrowhead and red arrow) but appeared decreased in more posterior clones (blue arrow). Examination of 9 clones in 6 discs from 2 independent experiments showed consistent changes. Images are partial projections for representative clone regions, Scale bar: 5 μm.

We next compared *pntP2* transcript levels in wildtype versus *pnt^p2^* null mutant retinal tissue. In both whole mutant eye discs (Fig. 7A-D) and in null mutant clones (Fig. 7E), increased *pntP2* transcription was evident in the MF and in the adjacent region where *pntP2* levels normally peak (Fig. 7A and S6). We also noticed a change not predicted by the RT-qPCR analysis, namely a decrease in *pntP2* transcripts in the posterior of the disc (Fig. 7A-D and S6). This suggests *pntP2* transcription is regulated differently in anterior versus posterior regions of the developing eye field. Reconsidering the RT-qPCR results in light of the spatial complexity to *pntP2* auto-regulation suggests that assay underestimated the fold increase in *pntP2* transcription that occurs in the anterior region where first round photoreceptor fates are specified.

Because PntP3 expression is strongest anteriorly (Fig. 2), we wondered whether the increase in *pntP2* transcripts detected in *pnt^p2^* mutant tissue might depend on cross-regulatory activation by PntP3. To test this we examined *pntP2* transcript levels in *pnt^p2p3^* double mutant clones (Fig. 7F). No increase was detected at the MF or in the adjacent region of peak expression. In more posterior *pnt^p2p3^* mutant clones, *pntP2* transcript levels were lower than in adjacent wild type clones, exactly as seen in *pnt^p2^* single mutant clones (Fig. 7E). Thus in anterior regions where PntP3 expression is strong, loss of PntP2 results in a PntP3-dependent increase in *pntP2* transcription whereas in posterior regions where PntP3 expression is normally low, loss of PntP2 results in a PntP3-independent reduction in *pntP2* transcription. As discussed below (Figure 8), we suggest that the regulatory relationships between Pnt isoforms established at the MF are reset after the SMW, and that the two different network topologies contribute specificity and robustness to first versus second round photoreceptor fate specification.

**Fig. 8.**
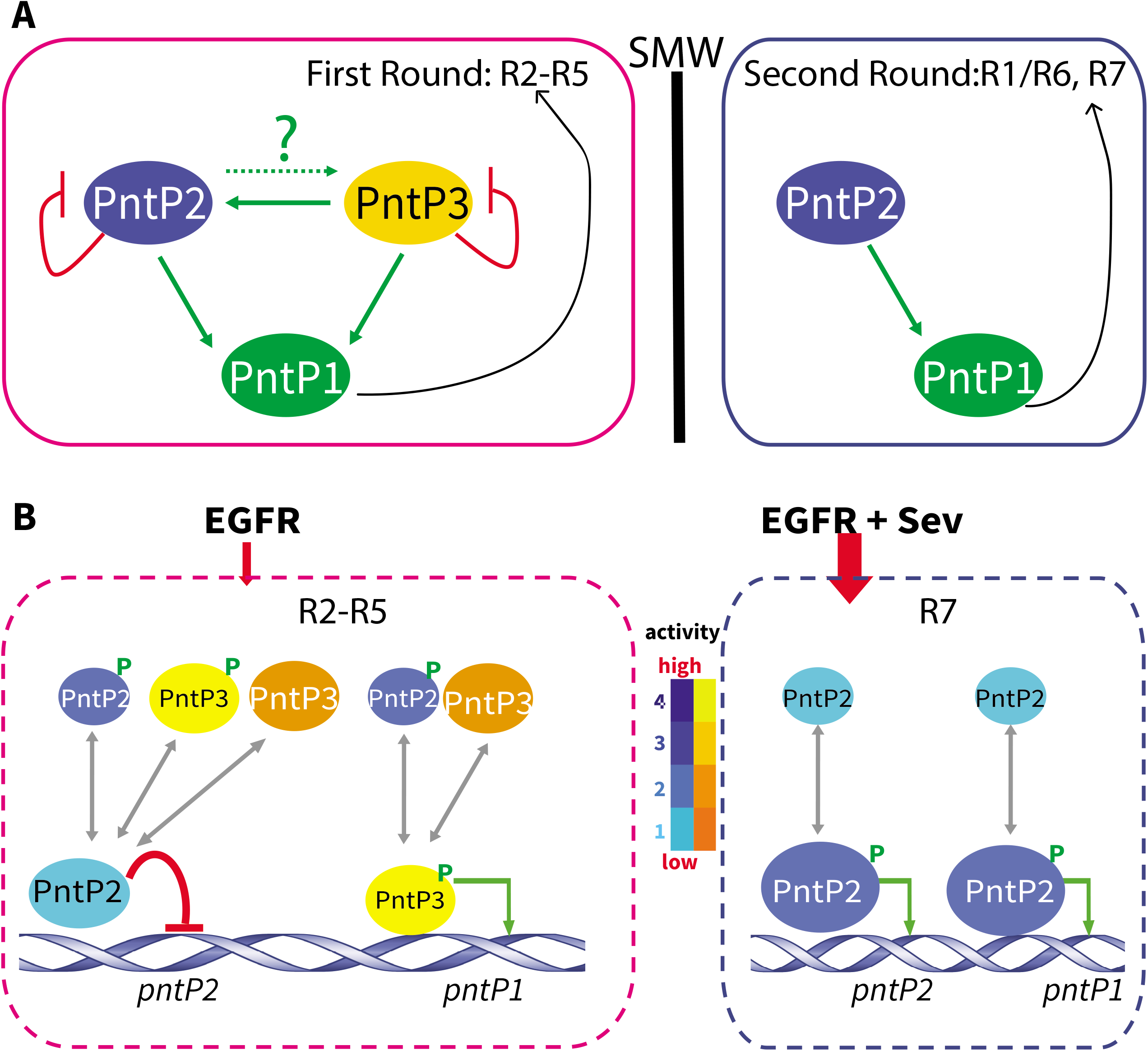
Model: Context-specific topology and function of the Pnt transcriptional regulatory network. **(A)** A schematic summary of the Pnt network. During first-round specification anterior to the SMW, both PntP2 and PntP3 auto-repress their transcription, PntP3 activates *pntP2*, and PntP2 and PntP3 redundantly activate *pntP1*. During the second-round, PntP2 auto-activates its own transcription and activates *pntP1*; PntP3 does not contribute. **(B)** Proposed network functions in the different signaling environments of first and second round specifications. The color scheme illustrates the range of transactivation activity: for PntP3, yellow indicates high activity and orange low; for PntP2 dark blue indicates high activity and light blue low. Different sized ovals depict relative abundance of the phosphorylated versus unphosphorylated forms. During first-round specification where there is only EGFR signaling, MAPK substrate competition between PntP2 and PntP3 keeps the ratio of phosphorylated to unphosphorylated protein low, reducing overall transactivation potential in the system. At the molecular level, if the enhancer of *pntP2* is biased towards unphosphorylated PntP2, this would limit access to the more active forms (PntP2p, PntP3 and PntP3p), effectively auto-repressing. The pool of more active forms would also be more likely to activate *pntP1* enhancer, even more so if it had a bias toward the phosphorylated forms. In contrast, regulation during the second round is simpler, with stronger MAPK activation from EGFR plus Sev signaling plus reduced substrate competition from PntP3 increasing the ratio of phosphorylated to unphosphorylated PntP2 to ensure robust activation of its own transcription and of *pntP1*.

## Discussion

In this study we explore the contributions of a previously uncharacterized Pointed isoform, PntP3, to the transcriptional effector network that directs developmental transitions downstream of receptor tyrosine kinase signaling. We show that PntP3, like PntP2, functions as a MAPK responsive transcription factor, but that despite their molecular and functional similarities, PntP3 and PntP2 have distinct expression patterns, transcriptional activities and mutant phenotypes. Together our results suggest that essential regulatory responsibilities previously attributed solely to PntP2, are actually distributed between PntP2 and PntP3, and that depending on context, the two work redundantly, uniquely or synergistically. We speculate that the network of auto- and cross-regulatory interactions we have uncovered between the isoforms fine-tunes Pnt transcriptional output to confer specificity and robustness to the developmental transitions it directs.

Our investigation of the PntP3 isoform has uncovered an unexpected bifurcation in the transcriptional effector network that transduces RTK/MAPK signaling. In doing so, it has also corrected an erroneous assumption regarding the role of the closely related PntP2 isoform. Prior to our study, the accepted model was that MAPK phosphorylation of PntP2, followed by PntP2p-mediated induction of *pntP1* transcription, provided the essential activating input for RTK-dependent transitions (Fig. 1C; Shwartz et al., 2013). As exemplified by studies in the eye, the genetic cornerstone of this model was that null alleles of either *pntP2* or *pntP1* produce identical phenotypes, namely a failure to specify photoreceptor R1-R7 fates (O’Neill et al., 1994; Shwartz et al., 2013; Yang and Baker, 2003). However the allele *pnt^Δ78^* (O’Neill et al., 1994), previously misinterpreted as a *pntP2*-specific null, actually disrupts the exon common to *pntP2* and *pntP3*. Thus the failure to specify R1-R7 fates reflects the compound loss of both PntP2 and PntP3. It should also be noted that an earlier study using hypomorphic truly *pntP2*-specific alleles concluded correctly that there is an “absolute requirement for *pntP2* function in R1, R6 and R7” but did not find a requirement in R2-R5 (Brunner et al., 1994a).

A schematic summarizing the combined contributions of PntP3 and PntP2 to photoreceptor fate specification is presented in Figure 8 as a framework for considering some of the mechanistic implications of our work. To recap briefly the key phenotypes and regulatory interactions on which the revised model is based, our study revealed redundant functional requirements for PntP2 and PntP3 in specifying the first round fates R2/R5/R3/R4; thus either single mutant recruits wild type 5-cell ommatidial clusters, while only in the double mutant are R2-R5 fates lost. Molecularly, PntP2 and PntP3 redundantly activate *pntP1* transcription (Fig. 8A) with significant reduction in *pntP1* levels detected only in the double mutant. In contrast, only PntP2 is required for R1, R6, R7 fates during the second round of photoreceptor specification and so eyes from isoform-specific *pnt^p2^* null mutants lack these three photoreceptors whereas *pnt^p3^* mutant ommatidia have the wild type complement of eight. Because *pntP1* transcript levels posterior to the SMW are already quite low in wild type discs, our FISH experiments were unable to detect the presumed reduction in *pntP1* in *pnt^p2^* mutant discs.

As a general developmental strategy, the redundant use of PntP2 and PntP3 may provide an effective buffer against genetic perturbations that reduce RTK signaling. Using R2-R5 photoreceptor specification as a specific example, the presence of redundant MAPK effectors in the early stages of ommatidial assembly may maximize overall robustness by minimizing early “mistakes” that would derail the entire process. Supporting this idea, we found that in a genetically sensitized background with reduced MAPK signaling output in R3, R4 precursors (Rebay and Rubin, 1995; Rebay et al., 2000), loss or reduction in dose of either *pntP2* or *pntP3*, which in otherwise wild type discs did not compromise patterning, now resulted in loss of these cell fates. Analogous results were obtained in the wing, suggesting that redundant use of PntP2 and PntP3 can confer robustness to genetic disruptions that reduce signaling. Given the broad participation of Pnt in specifying diverse cell fates throughout development, we anticipate future studies will uncover additional examples of context-specific redundant and unique functional requirements for Pnt isoforms.

Just as inadequate signaling compromises developmental transitions, so will excessive, oncogenic-levels of induction of transcriptional programs. For example, genetic perturbations that enhance RTK pathway output, such as increased Pnt expression or activity, severely disrupt ommatidial assembly and wing patterning (Brunner et al., 1994b; Karim et al., 1996; Prober and Edgar, 2000). Therefore to prevent redundant use of PntP2 and PntP3 from overactivating transcriptional programs, there need to be mechanisms to fine-tune and limit output.

The unexpected negative auto-regulation of both *pntP2* and *pntP3* transcript levels uncovered in our study may serve this purpose (Fig. 8A). Thus as detected by RT-qPCR for both *pntP2* and *pntP3*, and confirmed by FISH anterior to the SMW for *pntP2*, transcript levels of each isoform increased in the absence of the encoded protein product. Based on our prior work showing extensive Pnt chromatin occupancy across the *pnt* locus (Webber et al., 2018), we favor a mechanism in which direct auto-repression keeps PntP2 and PntP3 levels in check; however an indirect mechanism involving Pnt-mediated transcriptional activation of a repressive factor is equally plausible. If direct auto-regulation is used, the ability of Pnt to recruit and co-occupy enhancers with the ETS family repressor Yan and the corepressor Groucho uncovered in a recent study (Webber et al., 2018) could provide the repressive mechanism.

Counteracting the negative auto-regulation at *pntP2* and *pntP3*, we also uncovered positive transcriptional cross-regulation whereby PntP3 can activate *pntP2*. Thus in *pnt^p2p3^* double mutant clones, the increase in *pntP2* transcript levels that occurs in *pnt^p2^* single mutants was no longer observed. Again, we favor the simplest model of direct activation of *pntP2* by PntP3 (Fig. 8A), but cannot rule out more complicated indirect regulatory relays. Whether the converse cross-regulation of *pntP3* transcription by PntP2 occurs, and whether PntP2 and/or PntP3 positively auto-regulate their transcription anterior to the SMW remains to be assessed. However the decrease in *pntP2* transcript levels measured posterior to the SMW in *pnt^p2^* mutant tissue argues that positive auto-regulation is possible, making it plausible that such regulation could also help fine-tune PntP2/P3 levels and output during specification of first round fates.

How specific PntP2:PntP3 ratios influence the acquisition of different photoreceptor cell fates will be in an interesting focus for future work. Numerous studies have shown that regulatory networks can either amplify or suppress both the intrinsic noise (i.e. the randomness associated with mRNA/protein expression and degradation) and extrinsic noise (i.e. the variability caused by fluctuations in cellular processes or environment) of protein levels to influence cell fate decisions (Chang et al., 2008; Singh and Hespanha, 2009; Voliotis and Bowsher, 2012). Very speculatively, perhaps the network of auto-repressive and cross-activating interactions between PntP2 and PntP3 also tunes the cell-to-cell variation in Pnt isoform or Pnt target gene expression, thereby influencing the response to inductive signaling.

Another intriguing feature of the network of transcriptional interactions uncovered in our study is that corresponding to the switch from redundancy between PntP2 and PntP3 to the uniqueness of PntP2, the balance of PntP2 autoregulation shifts from repression during first round fate specification to activation during the second round (Fig. 8A). Fig. 8B offers speculation on how the distinct RTK signaling environments anterior vs. posterior to the SMW, combined with intrinsic differences in PntP2 vs. PntP3 activity, could produce this shift. Briefly, R1-R7 fates all rely on EGFR signaling while the R1, R6, R7 photoreceptors specified during the second-round experience additional RTK signaling through Sevenless (Sev); studies focused on R7 specification have highlighted the requirement for both EGFR and Sev (Basler and Hafen, 1989; Stark et al., 1976; Tomlinson et al., 2019). Use of the same Ras/MAPK/Pnt pathway means that EGFR and Sev-initiated signals can be considered interchangeable (Fortini et al., 1992; Freeman, 1996), with lower pathway activity required in the first round and higher activity needed in the second (Tomlinson et al., 2011; Tomlinson et al., 2019).

Because both PntP2 and PntP3 are direct MAPK substrates whose transactivation potential is increased by phosphorylation, their combined transcriptional output will be sensitive to the abundance of activated MAPK. Under conditions of lower pathway signaling and when both isoforms are co-expressed, as occurs anterior to the SMW, competition for the limited pool of activated MAPK will lead to domination by the unphosphorylated, less active forms. The presence of PntP3, whose unphosphorylated form has equivalent activity to PntP2p, and whose phosphorylated form has twice the activity of PntP2p (Figure 1D), may be important to make sure pathway output remains above a certain threshold in situations with lower levels of signaling. Although at first glance this might predict that the system would not tolerate loss of PntP3, because loss of PntP3 also reduces MAPK substrate competition, this would shift the distribution of PntP2 protein toward the phosphorylated more active form, thereby ensuring a robust transcriptional response.

How might these relationships manifest at the level of target gene enhancers? Given that PntP2 and PntP3 have the same ETS DNA binding domain and are identical except for the sequences N-terminal to the SAM, we expect they recognize the same DNA binding sites. Thus in the simplest scenario in which the phosphorylated and unphosphorylated forms of both PntP2 and PntP3 compete equally for enhancer occupancy, situations in which the unphosphorylated forms predominate would prevent excess activation of target genes. Much greater regulatory complexity is possible if modest enhancer-specific preferences between PntP2 and PntP3 and between the phosphorylated and unphosphorylated forms bias the competition. We suggest such biased competition will be essential to achieving limited activation, or even repression, of target genes such as *pntP2*, while allowing strong induction of others, such as *pntP1*, in the same cell. Based on a large-scale interactome study that reported closely related isoform pairs often have distinct protein-protein interaction patterns (Yang et al., 2016), it is possible that association with distinct cofactors also contributes to Pnt isoform enhancer occupancy bias. Much more complicated patterns of competition and cooperativity in which different Pnt isoforms and species co-occupy enhancers with each other and with heterologous transcription factors will undoubtedly contribute to the target gene-specific regulation needed to induce different cell types.

The substrate competition-based model also readily explains the transcriptional shifts that may occur in the individual *pnt^p2^* and *pnt^p3^* mutants. If one removes either PntP2 or PntP3, then overall competition for activated MAPK is eased, resulting in domination by the phosphorylated form of the remaining protein to boost transcriptional output. This would derepress targets like *pntP2*, as detected in our experiments, while activation of targets like *pntP1* would continue at physiologically functional levels. This same scenario plays out in an even stronger form in the wild type disc during specification of second round photoreceptor fates, where the combination of only PntP2 plus twice the RTK pathway input would result in phosphorylation of an even greater proportion of total PntP2 protein (Fig. 8B). Because PntP2 appears to have intrinsically weaker transaction potential than PntP3, ensuring full phosphorylation in situations where it is the sole MAPK effector may be critical to activating the transcriptional program.

Our study adds to the growing appreciation of the enormous and exquisitely sensitive regulatory potential available to developing tissues through the combinatorial expression and use of different protein isoforms, and also offers insights beyond the *Drosophila* arena. The human homologs ETS1 and ETS2 show intriguing structural and functional parallels to *Drosophila* PntP2 and PntP3 (Wasylyk et al., 1997; Watson et al., 1988). In particular, ETS1 and ETS2 have distinct N-termini, similar but not identical MAPK responsiveness, identical DNA binding domains, and overlapping but not identical functions and expression patterns. For both ETS1/2 and the PntP2/P3 isoforms, their distinct transcriptional activities are associated with the unique N-termini. Thus PntP2 and ETS2 both have longer N-terminal stretches and weaker transactivation activity than PntP3 and ETS1(Wasylyk et al., 1997). ETS2 is also able to switch between activating and repressing functions in a cell type specific manner (A Fry and Inoue, 2018), just as we have proposed for PntP2. Given the striking parallels between the fly and human contexts, continued exploration of the molecular mechanisms underlying Pnt-mediated transcriptional responses may provide new insight on signaling robustness and specificity in mammalian systems and related human diseases.

## Acknowledgements

We thank Jemma Webber and Matt Hope for initial analysis of PntP3 transcriptional activities; Matt Hope for construction of UAS and pMT constructs; Wei Du for *ey-FLP; act-Gal4, UAS-GFP/CyO; FRT82B, tub-GAL80/TM6B* flies; Jiajie Xu, Will Yee and Hitoshi Matakatsu in Rick Fehon’s lab for advice on RT-qPCR and CRISPR; Rachael Bakker in Rich Carthew’s lab and Jiacheng Zhang in Jingyi Fei’s lab for advice on FISH; Nicolas Pelaez in Rich Carthew’s lab for advice on GFP-PntP3 quantification; Claude Desplan for Salm antibody; Hugo Bellen for Sens antibody; Rick Fehon for the equipment to photograph adult eyes; Ed Munro, Chip Ferguson, Mike Rust, Rich Carthew, current and former members of the Rebay and Fehon labs including Trevor Davis, Nicelio Sanchez-Luege, Xiao Sun, Julio Miranda-Alban, and Suzy Hur for helpful discussions; and Chip Ferguson, Matt Hope and Jemma Webber for helpful comments on the manuscript. The work was supported by NIH R01 GM080372 to IR. Further support from NIH R01 EY025957 to IR and from the Genomics Core Facility through a University of Chicago Cancer Center Support Grant P30 CA014599 is acknowledged.

